# A Cdk1 phosphomimic mutant of MCAK impairs microtubule end recognition

**DOI:** 10.1101/188425

**Authors:** Hannah R. Belsham, Claire T. Friel

**Author notes:** Corresponding author: Claire Friel.

## Abstract

The microtubule depolymerising kinesin-13, MCAK, is phosphorylated at residue T537 by Cdk1. This is the only known phosphorylation site within MCAK’s motor domain. To understand the impact of phosphorylation by Cdk1 microtubule depolymerisation activity, we have investigated the molecular mechanism of the phosphomimic mutant T537E. This mutant significantly impairs microtubule depolymerisation activity and when transfected into cells causes metaphase arrest and misaligned chromosomes. We show that the molecular mechanism underlying the reduced depolymerisation activity of this phosphomimic mutant is an inability to recognise the microtubule end. The microtubule-end residence time is reduced relative to wild-type MCAK, whereas the lattice residence time is unchanged by the phosphomimic mutation. Further, the microtubule-end specific stimulation of ADP dissociation, characteristic of MCAK, is abolished by this mutation. Our data shows that T537E is unable to distinguish between the microtubule end and the microtubule lattice.

## Introduction

Mitotic Centromere Associated Kinesin (MCAK) is a member of the kinesin-13 family of microtubule depolymerising kinesins. MCAK plays crucial roles in the cell cycle both in building the mitotic spindle and in correcting erroneous microtubule-kinetochore attachments. Therefore, both the localisation and depolymerisation activity of MCAK must be tightly regulated throughout the cell cycle. This regulation is primarily achieved through phosphorylation.

MCAK is regulated through the action of various mitotic kinases, including the aurora kinases, polo like kinase 1 and p21 activated kinase 1(Andrews et al. 2004; Lan et al. 2004; Pakala et al. 2012; Zhang et al. 2011; Zhang et al. 2008). These kinases phosphorylate MCAK at various sites within the N and C-terminal domains and in the neck region. Only one phosphorylation site within MCAK’s motor domain has been identified to date. Threonine 537, which is located adjacent to the α4 helix on the tubulin-binding face of the motor domain (Figure 1A), is phosphorylated by the cyclin-dependant kinase Cdk1 (Sanhaji et al. 2010). A phosphomimic mutant at this position, T537E, has reduced depolymerisation activity and overexpression of this mutant in cells leads to misaligned chromosomes, metaphase arrest and reduced intercentromeric distances (Sanhaji et al. 2010).

**Figure 1.**
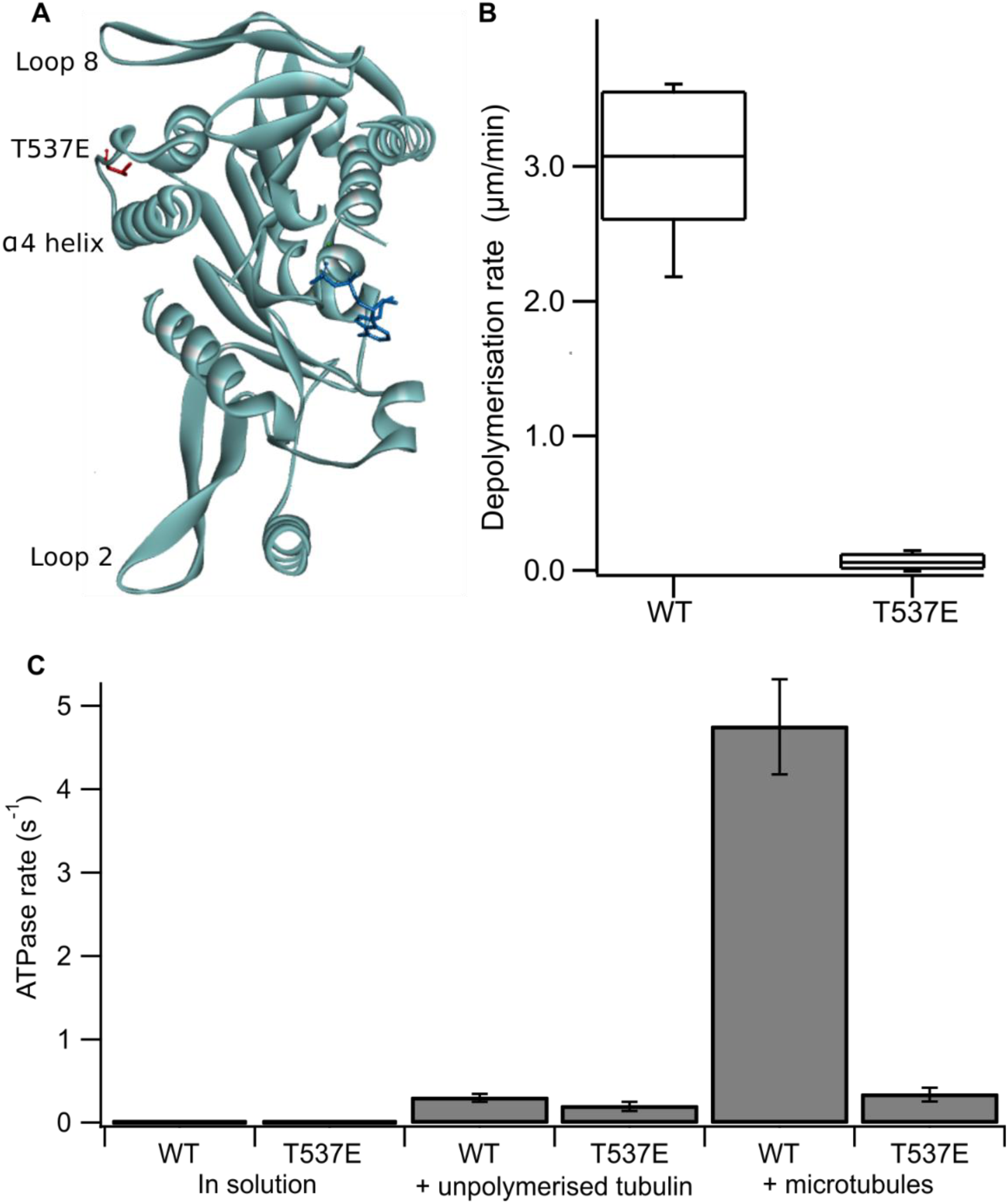
The phosphomimic mutant T537E has reduced depolymerisation activity and reduced microtubule-stimulated ATPase activity. (A) Location of T537 in the crystal structure of the MCAK motor domain (PDB ID: 2HEH). T537 (red) is located adjacent to the α4 helix. The α4 helix, loop 2 and loop 8 are at the microtubule binding interface of MCAK. Nucleotide is shown in dark blue (B) Depolymerisation rate of WT MCAK and T537E mutant. The distribution of the rates of depolymerisation of single microtubules after the addition of 40 nM MCAK is shown in the box plots. (C) ATPase rate of WT MCAK and T537E mutant in solution, in the presence of 10 µM unpolymerised tubulin and in the presence of 10 µM microtubules.

MCAK is a specialist microtubule depolymerising kinesin and its ATP turnover cycle is tailored to this function. While many kinesins have a translocating, “walking” action along the microtubule, MCAK diffuses on the microtubule lattice in the ADP-bound state. The microtubule end specifically accelerates ADP dissociation, stimulating exchange of ADP for ATP, which allows MCAK to bind tightly at the end of the microtubule. The binding of ATP.MCAK at the microtubule end promotes dissociation of tubulin dimers and thereby microtubule depolymerisation (Friel & Howard 2011).

To understand how phosphorylation by Cdk1 impairs MCAK’s depolymerisation activity we studied the effect of the phosphomimic substitution T537E on the molecular mechanism of microtubule depolymerisation. Here we show that the phosphomimic mutant, T537E, cannot distinguish between the microtubule end and the microtubule lattice, an ability characteristic of wild-type MCAK and other kinesins which regulate microtubule dynamics.

## Results

### The phosphomimic mutant T537E reduces depolymerisation activity and microtubule-stimulated ATPase activity

Firstly, we confirmed the effect of the substitution T537E on MCAK’s depolymerisation activity *in vitro*. It has been shown previously at high concentration (500 nM) that the mutation T537E decreases depolymerisation activity 4-fold relative to wild-type MCAK. (Sanhaji et al. 2010). We measured depolymerisation activity at 40 nM, the concentration at which the fastest microtubule depolymerisation for wild-type MCAK is observed (Helenius et al. 2006). Under these conditions the phosphomimic mutant had a depolymerisation rate 50-fold slower than the wild-type (0.06 ± 0.06 µm min^-1^ and 3.04 ± 0.53 µm min^-1^ (mean ± standard deviation), respectively) (Figure 1B). Thus, confirming that the phosphomimic substitution dramatically decreases MCAK’s depolymerisation activity.

We next measured the ATPase activity of T537E in the absence of tubulin, in the presence of unpolymerized tubulin and in the presence of microtubules. The ATPase rate of the T537E mutant in solution was not significantly different to wild type (5.33 ± 0.33 x 10^-3^ s^-1^ compared with 4.47 x 10^-3^ ± 2.60 x 10^-3^ s^-1^, p = 0.6002)(Figure 1C).Similarly, the ATPase in the presence of unpolymerized tubulin was not significantly affected by the phosphomimic mutation: 0.194 ± 0.055 s^-1^ compared with 0.299 ± 0.047 s^-1^ for wild-type MCAK, p = 0.0658 (Figure 1C). These data indicate that the phosphomimic mutant folds correctly as it turns over ATP at a similar rate to wild-type both in solution and in the presence of unpolymerized tubulin. These data also suggest that T537E can still interact with unpolymerized tubulin, as its ATPase rate is accelerated by unpolymerized tubulin to the same degree as wild type. By contrast, there is a dramatic difference in the microtubule-stimulated ATPase activity of T537E relative to wild-type MCAK. The microtubule-stimulated ATPase rate of T537E is 14-fold slower than wild type MCAK: 0.335 ± 0.081 s^-1^ and 4.75 ± 0.055 s^-1^, respectively (Figure 1C).

### The mutation T537E abolishes long microtubule end residence events

The ATPase activity of wild-type MCAK is maximally accelerated by microtubule ends (Friel & Howard 2011; Hunter et al. 2003). The residue T537 is near the α4 helix of the MCAK motor domain (Figure 1A). Residues in the α4 helix are critical to MCAK’s ability to recognise the microtubule end (Patel et al. 2016). This proximity to the α4 helix together with the observation that the mutation T537E specifically impairs the microtubule stimulated ATPase of MCAK suggests that this phosphomimic mutation may also interfere with MCAK’s ability to distinguish the microtubule end from the lattice. We used TIRF microscopy to observe the interaction of single molecules of MCAKGFP and T537E-GFP with microtubules. Wild type MCAK makes short diffusive interaction with the microtubule lattice. However, when it reaches the microtubule end ADP dissociation is accelerated, leading to nucleotide exchange and ATP.MCAK binds tightly and displays longer microtubule end residence events (Figure 2A). By contrast, T537E does not show long end binding events (Figure 2A). Quantification of microtubule end residence times for wild-type MCAK and T537E shows that 32 % of molecules for the wild type but only 0.4% of molecules for T537E stayed at the microtubule end for longer than 2 s (Figure 2B). The affinity for the microtubule lattice is not significantly changed for this mutant. The lattice residence time for T537E is unchanged relative to wild type (0.42 ± 0.34 s and 0.48 ± 0.02 s, respectively) and neither the association or dissociation rates are significantly different (Supplementary Figure S1 and Supplementary Table S1). However, the end residence time is decreased to 0.64 ± 0.02 s from 2.03 ± 0.13 s for the wild type and the rate of dissociation from the microtubule end is increased relative to wild-type (Supplementary Table S1). These data imply that the molecular mechanism underlying the attenuation of depolymerisation activity due to this phosphomimic mutation is loss of ability to distinguish between the microtubule end and the microtubule lattice.

**Figure 2:**
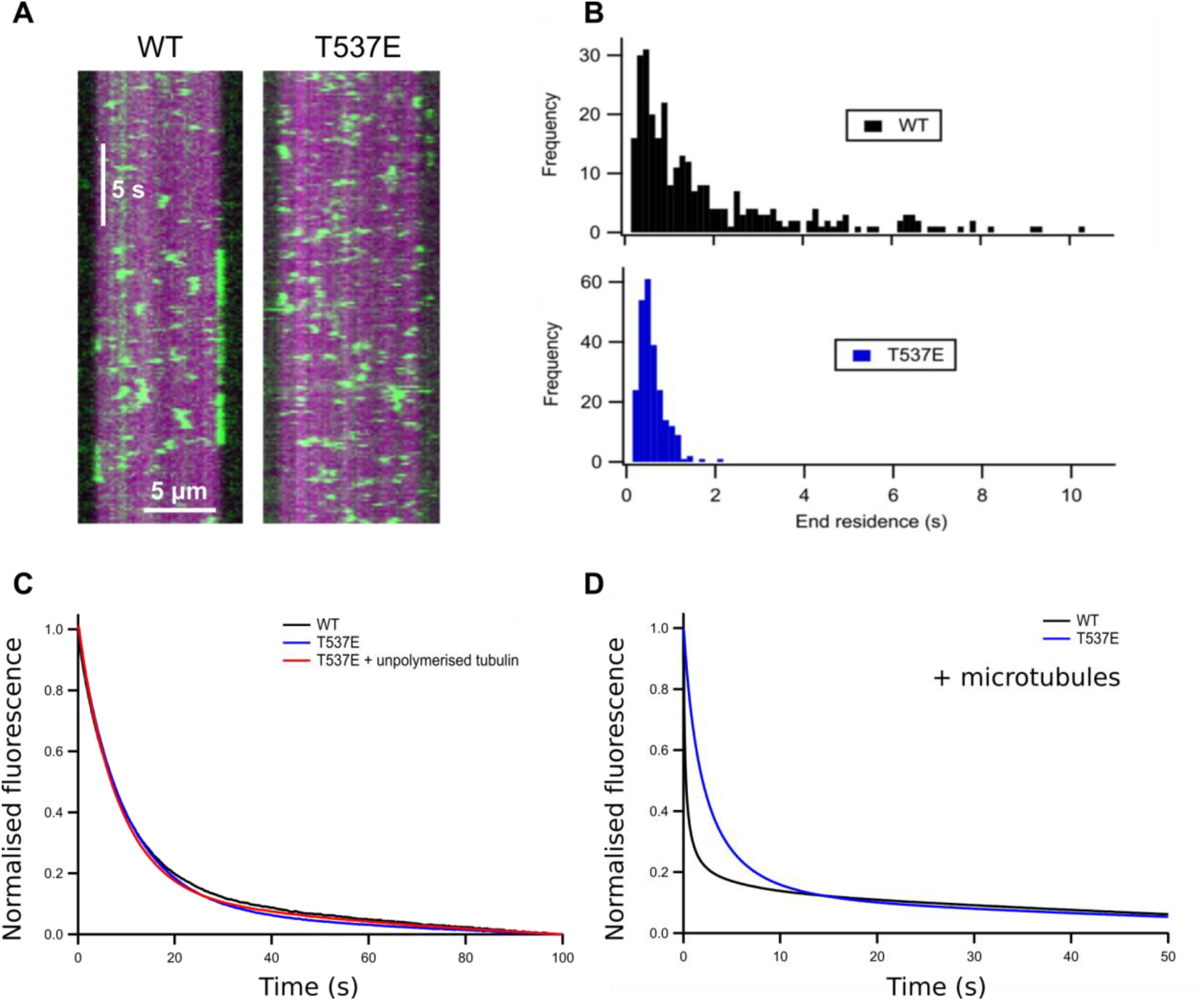
T537E has few long microtubule-end binding events and has lost microtubule-end stimulated mantADP dissociation. (A) Kymographs showing the interaction of WT MCAK and the T537E mutant with the microtubule. (B) Histograms showing the end residence time of single molecules of WT MCAK and T537E. Wild type data published previously (Patel et al. 2016). (C) Representative fluorescence traces for the dissociation of mantADP from WT MCAK and T537E in solution and for T537E in the presence of 10 µM unpolymerised tubulin. (D) Representative fluorescence traces for the dissociation of mantADP from WT MCAK and T537E in the presence of 5.7 µM microtubules.

### The microtubule end does not accelerate ADP dissociation from T537E

ATP turnover by MCAK is maximally accelerated by the microtubule end due to microtubule end specific acceleration of ADP dissociation. To test whether the difference in T537E interaction with the microtubule end had affected the ability of the microtubule end to accelerate ADP dissociation, we measured the rate of dissociation of ADP tagged with the small fluorophore mant (mantADP). In solution and in the presence of unpolymerised tubulin, the rate constant for ADP dissociation was not significantly affected by this mutation (WT 0.102 ± 0.013 s^-1^, T537E 0.114 ± 0.013 s^-1^, WT + Tb 0.114 ± 0.024 s^-1^, T537E + Tb 0.120 ± 0.019 s^-1^)(Table 1 and Figure 2C). This is in agreement with the ATPase activities under these conditions which are not significantly changed.

**Table 1.** Compiled results of depolymerisation rate, ATPase activity, microtubule-end residence time and mantADP dissociation rates for wild type MCAK and T537E. *The data and fits to the data from which these rate constants were obtained are shown in Supplementary Figure S2.

| MCAK variant |  | WT | T537E |
| --- | --- | --- | --- |
|  |  | **($\mu\text{m}/\text{min}$) (mean ± SD)** |  |
| **Depolymerisation Rate** | | $3.04 \pm 0.53$<br>(n = 20) | $0.06 \pm 0.06$<br>(n = 20) |
| **ATPase activity** |  | **($\text{s}^{-1}$) (mean ± SD)** |  |
| Solution (basal) | | $0.00447 \pm 0.0026$<br>(n = 3) | $0.00533 \pm 0.0033$<br>(n = 3) |
| Tubulin-stimulated | | $0.299 \pm 0.047$<br>(n = 3) | $0.194 \pm 0.055$<br>(n = 3) |
| Microtubule-stimulated | | $4.75 \pm 0.057$<br>(n = 3) | $0.335 \pm 0.081$<br>(n = 3) |
|  |  | **(s) (mean ± SEM)** |  |
| **Microtubule-end residence time** | | $2.03 \pm 0.13$<br>(n = 238) | $0.64 \pm 0.02$<br>(n = 276) |
| **mantADP dissociation\*** |  | **($\text{s}^{-1}$) (mean ± SD)** |  |
| Solution | | $0.102 \pm 0.013$<br>(n = 3) | $0.114 \pm 0.013$<br>(n = 3) |
| Tubulin-stimulated | | $0.114 \pm 0.023$<br>(n = 3) | $0.120 \pm 0.019$<br>(n = 3) |
| Microtubule-stimulated | First phase | $4.11 \pm 0.41$<br>(n = 3) | $0.369 \pm 0.089$<br>(n = 3) |
| | Second phase | $0.341 \pm 0.051$<br>(n = 3) | n/a |

In the presence of microtubules, the change in fluorescence associated with mantADP dissociation from wild type MCAK is best described by a double exponential function. The two phases can be explained as a microtubule-end stimulated fast phase and a slower phase corresponding to molecules which do not encounter the microtubule end (*i.e.* remain in solution, in contact with unpolymerised tubulin or in contact with the microtubule lattice over the course of the experiment). The fast phase of mantADP dissociation from wild type MCAK measured here has a rate constant of 4.11 ± 0.41 s^-1^ and the slower phase a rate constant of 0.341 ± 0.051 s^-1^. The rate constant for the slower phase is in close agreement with the rate constant for mantADP dissociation from MCAK in solution or in the presence of unpolymerized tubulin. This is in agreement with this phase representing mantADP dissociation from molecules which do not contact the microtubule end. By contrast with WT-MCAK, the change in fluorescence associated with mantADP dissociation from T537E was well described by a single exponential function with a rate constant of 0.369 ± 0.090 s^-1^. The faster phase was lost for the phosphomimic mutant indicating that the microtubule end cannot accelerate mantADP dissociation from T537E. In fact, the rate constant for the single phase observed for mantADP dissociation from T537E is not significantly different from the slower phase of mantADP dissociation from wild type MCAK (p = 0.6679, Figure 2D and Table 1). This observation, that the single kinetic phase observed for T537E has a rate constant in agreement with the kinetic phase for WT-MCAK that represents molecules that do not reach the microtubule end, indicates that the phosphomimic mutant T537E has lost the ability to distinguish between the microtubule lattice and the microtubule end.

## Discussion

To understand the molecular mechanism by which phosphorylation by Cdk1 impairs MCAK’s depolymerisation activity, we studied the phosphomimic mutant T537E. This mutation causes a 50-fold decrease in the rate of microtubule depolymerisation by MCAK. Overexpression of this phosphomimic mutant in cells shows that it can localise to centromeres but that chromosome alignment is disrupted and cells arrest in metaphase (Sanhaji et al. 2010). The intra-centromere distance is decreased suggesting that cells expressing this mutant form of MCAK cannot generate tension across centromeres or correct erroneous chromosome attachments.

We have shown that, whilst the in solution and unpolymerized tubulin-stimulated ATPase activity of the phosphomimic mutant are unchanged, the microtubule stimulated ATPase is reduced 14-fold relative to wild-type MCAK. Single molecule TIRF analysis of this mutant showed that this impact on the microtubule-stimulated ATPase was due to loss of the ability of T537E to recognise the microtubule end. We saw that long end residence times, characteristic of wild-type MCAK’s interaction with the microtubule, are abolished by the T537E mutation. Further, T537E cannot undergo microtubule-end stimulated acceleration of ADP dissociation, a crucial requirement for the depolymerisation activity of MCAK. Together our data show that T537E, unlike wild-type MCAK, cannot distinguish between the microtubule lattice and the microtubule end.

The residue T537, the target of phosphorylation by Cdk1 and the only currently identified phosphorylation site within the MCAK motor domain, is located adjacent to the C-terminal end of the α4 helix. The α4 helix has been suggested, on the basis of currently available structures of Kinesin-13 motor domains (Asenjo et al. 2013; Ogawa et al. 2004; Shipley et al. 2004; Wang et al. 2017), to play a role in deforming tubulin dimers thereby destabilising the microtubule and promoting depolymerisation. The ATP turnover cycle of MCAK differs from translocating kinesins and is adapted to promote tight binding of MCAK at the microtubule end where it can have maximal depolymerising impact (Friel & Howard 2011). Previous work from our lab has shown that mutations in the α4 helix impact the molecular mechanism of MCAK in the same way as the phosphomimic mutant described here (Patel et al. 2016). The proximity of T537 to the α4 helix may also provide the key to why the T537E mutant is unable to distinguish between the microtubule lattice and the microtubule end. The α4 helix is proposed to have a larger interface with tubulin at the intradimer groove in a curved conformation of tubulin, as may be found at the microtubule end, compared with a more constrained straight conformation within the microtubule lattice (Asenjo et al. 2013; Ogawa et al. 2004; Wang et al. 2017). Disruption of this crucial interface by mutation or by phosphorylation impairs MCAK’s ability to distinguish different tubulin conformations and thereby recognise the microtubule end.

Our data shows that the T537E mutant, although significantly impaired in depolymerisation activity, can interact with the microtubule lattice with an affinity similar to wild-type MCAK. This mutant still displays the characteristic diffusive interaction of MCAK with the microtubule lattice and can still reach the microtubule end. Assuming that the behaviour we have observed is a good representation of MCAK’s activity in cells, this would allow MCAK phosphorylated at T537 to interact with microtubules of the mitotic spindle and localise at microtubule ends close to centromeres but locked in an inactive state that is blocked from recognising the microtubule end and promoting depolymerisation. Phosphorylation by Cdk1 holds MCAK in the set position; dephosphorylation is then the starter’s gun allowing MCAK to instantly begin depolymerising kinetochore-attached microtubules at the correct moment. Phosphorylation at the Cdk1 site, T537, rather than blocking MCAK’s interaction with microtubules, holds MCAK in an inactive state whilst allowing correct localisation. Thus, permitting the rapid switching of activity characteristic of regulation by phosphorylation.

## Methods

### Proteins

Full length human MCAK-his6 and MCAK-his6-EGFP in wild type and mutated forms were expressed in Sf9 cells and purified using nickel affinity chromatography as described previously (Helenius et al. 2006). MCAK concentrations are given as monomer concentrations. Porcine brain tubulin was purified as described previously (Castoldi & Popov 2003). Single cycled, fluorescently labelled microtubules and double cycled microtubules were prepared as described previously (Patel et al. 2016). The concentration of microtubules is given as the concentration of polymerised tubulin.

### Microtubule depolymerisation

Microtubule depolymerisation rates were determined by measuring the length of immobilised, GMPCPP-stabilised, rhodamine labelled, single cycled microtubules over time after the addition of 40 nM MCAK as described previously (Patel et al. 2016).

### ATPase assays

ATPase rates in solution were measured using 3 µM MCAK and monitoring the production of ADP by HPLC as described previously (Friel et al. 2011; Friel & Howard 2011; Patel et al. 2016). For assays with tubulin or double-cycled microtubules 0.1 µM MCAK was used and the production of ADP was monitored by linking it to the oxidation of NADH (De La Cruz & Ostap 2009). For both assays the change in concentration of ADP per second was divided by the concentration of MCAK to give the ATPase activity per second per motor domain.

### Single molecule TIRF assays

Single molecules of MCAK-GFP were observed on immobilised, GMPCPP-stabilised, rhodamine labelled, single cycled, microtubules using TIRF microscopy as described previously (Patel et al. 2016). Kymographs for individual microtubules were used to measure the time individual MCAK-GFP molecules spent at the microtubule end and on the lattice.

### ADP dissociation assays

The dissociation of mantADP from MCAK was measured as described previously (Patel et al. 2014), in solution or with the addition of 10 µM tubulin or 5.7 µM double cycled microtubules (chosen to have a comparable number of microtubule ends as the ATPase assay with microtubules). The fluorescence traces were fitted to a single exponential, or double exponential if required, with an additional linear function to account for the photobleaching of mant.

## Acknowledgements

We are grateful to Alex Rathbone for technical support. Microscopy assays were carried out in the University of Nottingham School of Life Sciences Imaging (SLIM) facility. We thank the SLIM team and in particular Chris Gell for supporting our work.

